# Seasonality but not photoperiodism affects susceptibility of the two-spotted spider mite, *Tetranychus urticae* Koch (Acari: Tetranychidae) to pesticides

**DOI:** 10.1101/2022.03.09.483610

**Authors:** Zhenguo Yang, Zinan Wang, Jing Ni, Aisi Da, Daoyan Xie, Henry Chung, Yanjie Luo

**Affiliations:** Sericulture and Apiculture Research Institute, Yunnan Academy of Agricultural Sciences, Yunnan, China; Key Laboratory of Green Prevention and Control of Agricultural Transboundary Pests of Yunnan Province, Yunnan Academy of Agricultural Sciences, Yunnan, China; Department of Entomology, Michigan State University, East Lansing, MI, USA; Ecology, Evolution, and Behavior Program, Michigan State University, East Lansing, MI, USA

## Abstract

Understanding how endogenous and exogenous factors such as annual seasonal rhythm and photoperiodism affect the toxicity of pesticides can help design integrated pest management strategies. The two-spotted spider mite *Tetranychus urticae* Koch (Acari: Tetranychidae), a worldwide phytophagous pest species distributed in areas with different time zones, is a good model to explore how the photoperiodism and seasonality affect the pesticide toxicity. In this study, we conducted a laboratory experiment from March 2017 to November 2018 where spider mites were reared at three photoperiod regimes in environmentally controlled incubators. The toxicities of two acaricides, propargite and diafenthiruon, were measured on *T. urticae* every month. To determine potential mechanisms underlying the changes in the acaricide toxicity, we measured body size and total GSTs activity with the toxicity measurements in 2018. Our results showed that the toxicities of the two acaricides were not significantly different among the three photoperiod regimes. However, both had a consistent trend along the season which increased in the spring and summer and decreased in the winter in all photoperiod regimes for two consecutive years, suggesting seasonality be an endogenous factor affecting the susceptibility of the spider mites to pesticide. Pearson’s correlation analyses showed only the body size had a weak negative correlation with the acaricide toxicities, suggesting a minor contribution to the higher toxicity from the smaller size of *T. urticae*. Our study is the first to show that seasonality can have an endogenous effect on the pesticide toxicity, and the results can inform practical insights into the pest management strategies.

## Introduction

Understanding the efficacy of chemical applications to pest species under different rhythmicity such as photoperiodism and seasonality can lead to better pest management strategies [1, 2] Biological rhythmicity is one of the most important systems that the arthropods use to coordinate their development and physiology[3]. Together with daily or seasonal alterations, changes in different abiotic factors such as photoperiod, temperature and humidity further shape the adaptations of arthropod behavior and physiology [4]. The cascading effects could indirectly influence the efficacy of pest control measures such as the use of pesticides in the field^2^. For example, the interaction of temperature and photoperiod has been found to induce diapause signals to prolong the nymphal development of two spider mite species, including *Tetranychus kanzawai* Kishida and *T. urticae* Koch (Acari: Tetranychidae) [5, 6], which further alters the capability of metabolizing multiple toxicants and pesticides[7]. Several studies reported that some insects and mite display circadian rhythm of susceptibility to toxic agent. For example, the toxicities of pesticides such as dimethyl 2, 2-dichlorovinyl phosphate (DDVP) [8], dicofol [9], methyl parathion [10], and permethrin [11] have been reported to correlate with the circadian rhythm in diverse pest species. This suggests that circadian rhythm be an important factor to be considered in pest management.

Seasonality is another rhythmal factor regulating arthropod physiology, potentially affecting the toxicity of pesticides. However, this is rarely studied under laboratory conditions. Current understanding about the seasonal effects comes from different physiological measures of field-collected organisms across the season, e.g., the seasonal susceptibility of honeybee worker (*Apis mellifera L.*) to several commonly used pesticides such as organophosphate, benzoylphenyl urea, carbamate, oxadiazine and pyrethroid [12, 13]. These studies provide a general understanding of the seasonal changes affecting the toxicity of pesticides. However, physiological adaptations across the seasons usually interact with multiple biotic and abiotic factors in the field, which could confound the analyses and results. It is less understood whether the arthropods have endogenous physiological adaptations to seasonality and whether these adaptations could affect pesticide toxicity. One way to investigate the endogenous effects of seasonality is to control abiotic factors such as temperature, humidity, and photoperiod in the laboratory during investigating the physiological changes and pesticide toxicity. Integrating these effects of rhythmal factors to pesticide toxicity into pest management has received much theoretical attention but less applied to pest controls in practical [1, 7, 14]. This could be due to difficulties in applying and controlling biological rhythms in the field. One of a few applications is on Lepidopteran storage pests, which regulated the timing and dosage of chemical applications based on circadian rhythm so as to minimize potential developments of insecticide resistance [15].

The two-spotted spider mite, *T. urticae*, is a worldwide phytophagous pest and has caused significant yield loss in multiple crops, including fruits, cotton, vegetables, and ornamentals [16, 18]. Due to their wide host range, small body size, short developmental time, and high fecundity, it has become one of the most serious pests to manage [18]. Chemical control has formed the foundation of tactics adopted by farmers to manage *T. urticae*, with propargite and diafenthiruon among the most effective and used acaricides to manage this pest [18]. A previous study showed no difference in both toxicities of propargite and diafenthiruon for *T. urticae* populations reared from three different photoperiod regimes in the summer [18]. However, whether the adaptations of *T. urticae* to different photoperiod regimes across the seasons affects the pesticide toxicity is unknown. To manage this mite pest more systematically, it is important to understand how the photoperiod and seasonality could affect the toxicities of commonly used pesticides in a relatively long time period.

In this study, we investigated the effects of the photoperiodism and seasonality on the toxicities of the two acaricides, propargite and diafenthiruon, to *T. urticae* in the laboratory. Our previous study suggested that photoperiodism does not affect acaricide toxicities in the summer [19]. However, it is unknown if the interaction between photoperiodism and seasonality could affect the toxicity. We hypothesized that photoperiod susceptibility to pesticide of mite has an intrinsic rhythm that adapts to the seasonal variation. To test this hypothesis, we reared *T. urticae* at three photoperiod regimes (10L:14D, 12L:12D, and 18L:6D) representing different exposure of light in three environmental incubators set with consistent temperature and humidity from March 2017 to November 2018. The temperature and humidity in incubators are controlled so that *T. urticae* do not enter diapause. To investigate the effects of photoperiodism and seasonality, toxicities of these two acaricides were measured every month for *T. urticae* in the incubators at the three photoperiod regimes. As body size can be treated as a proxy of fitness [20, 21] and the glutathione S-transferases (GSTs) is one of the detoxification enzyme super-family of which the toxicity of different pesticides were found under circadian control of this enzyme in several arthropods [22-24], we included the body size measurement and GSTs quantification along with our toxicity assays. Results from this study can provide a new perspective towards understanding relationships between the photoperiodism, seasonality, and pesticide toxicity in arthropods and provide inputs towards developing integrated pest management strategies.

## Materials and Methods

### Mite sources

All experiments and measurements in this study were conducted on laboratory populations of *T. urticae*, collected from rose tree plantation at Kunming, China (25.15°N, 102.76°E) in 2010. The populations were maintained on kidney bean leaves (*Phaseolus vulgaris* L.) in the climate-controlled incubators (GXZ-250A, Ningbo Jiangnan Instrument Co., Ltd., Zhejiang, China) set with a photoperiod of 14L:10D, a temperature of 25 ± 0.5°C, and a relative humidity of 75 ± 5%. White light-emitting diodes (LEDs) were used with light intensity being 500 – 700 Lx. In this study, three laboratory populations were established in 2015 at three photoperiod regimes, including 10L:14D, 12L:12D, and 18L:6D, with all other abiotic conditions kept constant. All assays and measurements were conducted on female adults sampled every month from March 2017 to November 2018. All populations reared in the incubators had not been exposed with any pesticide or acaricide.

### Toxicity bioassay

The Potter spray tower bioassay method [25]was used to determine the toxicity of propargite (90%, Yisheng Chemical Technology Co., Ltd., Shanghai, China) and diafenthiruon (98%, Qingdao Pesticide Factory, Shandong, China) on *T. urticae* populations under three photoperiod regimes every month from March to December, 2017 and January to November, 2018. Both diafenthiruon and propargite were diluted using double-distilled water (ddH_2_O) to five different concentrations, including 3.125, 6.25, 12.5, 25, and 50 μg/mL for diafenthiruon, and 62.5, 125, 250, 500, and 1000 μg/mL for propargite. The ddH_2_O was used as the control. Each treatment has three replicates. For each replicate, 25 – 30 females were transferred to a 50-mm-diameter bean leaf disk which was placed on a 90-mm-diameter Petri dish and soaked with wet cotton. All leaf discs and mites were maintained on dish for up to 4 h before the assay. After a 10-second settling time, a Potter spray tower (Burkard, Uxbridge, UK) which producing a deposit of 9.4 mg/cm^2^ at 25 ± 0.5°C was used to spray leaf discs. The sprayed leaf discs were maintained at same condition for 24 h until examining the mortality of mites. The movement of any appendage after gentle stimulation with a fine brush was used as the criterion to determine the mortality of the mites.

### Adult body size

Body size was measured at the same time of the GSTs activity bioassay every month from January to November, 2018. Following the method in Danielson-Francois et al. [21], in each month, three replicates of 50 females were randomly selected from the population under different photoperiod regimes and fixed on the slides using a double-sided tape. Sampled mites were placed under the light microscope (GL-99TI, Guiguang Instrument Co., Ltd. Guangxi, China). Body length and width was measured using the micromorphology instrument (Dongfang Agricultural Biotechnology Co., Ltd., Beijing, China). The size index (= body length × body width) was used to determine body size and used for statistical analyses.

### Glutathione S-transferases (GSTs) activity assay

To determine whether GSTs contribute to the difference of toxicity between populations and across the season, three replicates of 50 females were randomly selected from each population for assaying of GSTs activity. The sampled females were weighed and homogenized in a 1 L ice-cold sodium phosphate buffer (0.04 M, pH 6.5). The homogenate was centrifuged at 10,000 × g for 10 mins at 4°C, and the supernatant which contains the GST enzymes was transferred into a clean micro-centrifuge tube. Total mite GSTs activity was determined using a Glutathione S-transferase (GSTs) activity test kit (Micro method kit, Solarbio, Beijing, China) following the manufacturer’s instructions. The total amount of GST activity was determined as the absorbance at 340 nm using a spectrophotometer (xMarkTM Microplate Reader, BIO-RAD, USA) at 15 s interval during a five-min reaction period at 37°C.

### Statistical analyses

To determine the effects of photoperiodism and seasonality to the toxicity of propargite and diafenthiruon on *T. urticae* populations, the mortality data under toxicity assays of the two acaricides were respectively subjected to probit analyses (PROC PROBIT, SAS v9.4, SAS Institute Inc., Cary, NC, USA) for each year, with the dose, photoperiod, time, and the interaction between photoperiod and time being the fixed terms (= Dose + Photoperiod + Time + Photoperiod × Time). The 50% lethal concentration (LC_50_) with 95% confidence interval was used to represent the toxicity of each acaricide for each month. To determine the effects of photoperiodism and seasonality to the body size and overall GSTs activity, we applied a linear mixed model (PROC MIXED, SAS v9.4, SAS Institute Inc., Cary, NC, USA) to the body size index and total amount of GSTs activity with the photoperiod, time and the interaction being fixed terms (= Photoperiod + Time + Photoperiod × Time). Tukey’s HSD method was used for the post hoc analyses at alpha = 0.05. In addition, to further understand whether the body size and GSTs activity contribute to the toxicity of the two acaricides, the Pearson’s correlation coefficients were used to characterize the relationship with the confidence interval reported.

## Results

### Toxicities of diafenthiruon and propargite to T. urticae correlate with seasonality but not photoperiodism

To determine whether and how the toxicities of diafenthiruon and propargite were affected by the seasonal changes and photoperiod, probit analyses were applied to the mortality data of monthly toxicity bioassay with dose, photoperiod, time and the interaction of photoperiod and time for both years, respectively. In 2017, only the time term showed a significant effect on the toxicity of both diafenthiruon (χ(8) = 172.3, P < 0.0001, **Fig 1 a**) and propargite (χ(8) = 81.5, P < 0.0001, **Fig 1 b**). Similar effects from the time term were found in 2018 (diafenthiruon: χ(10) = 59.4, P < 0.0001, **Fig 1 c**, propargite: χ(10) = 58.6, P < 0.0001, Fig. 1 d). However, both the photoperiod and the interaction terms did not have any effects on the toxicities for both acaricides (**Fig 1 a-d**). In 2017, the susceptibility of *T. urticae* to both acaricides had same levels from March to September, started decreasing from October, and reached their minimum on December (**Fig 1**). For example, the LC_50_ of diafenthiruon on *T. urticae* at Photoperiod 12L:12D increased from 7.5 (95%CI: 5.0 – 11.5) mg/L on March to 17.1 (95%CI: 14.5 – 20.2) mg/L on December (**Fig 1 a**), and the LC_50_ of propargite at Photoperiod 12L:12D increased from 190.1 (95%CI: 159.3 – 227.1) mg/L on March to 442.4 (95%CI: 362.1 – 540.5) mg/L on December in 2017 (**Fig 1 b**). Similar changing trends of the toxicities across seasons were observed in 2018 that toxicities started decreasing from January to March, remained at similar levels for the next eight months. Results for both years showed that the toxicities of both acaricides to *T. urticae* were affected by annual seasonal variation but not photoperiodism in constant temperature (25 ± 0.5°C) and constant relative humidity (75 ± 5%).

**Fig 1.**
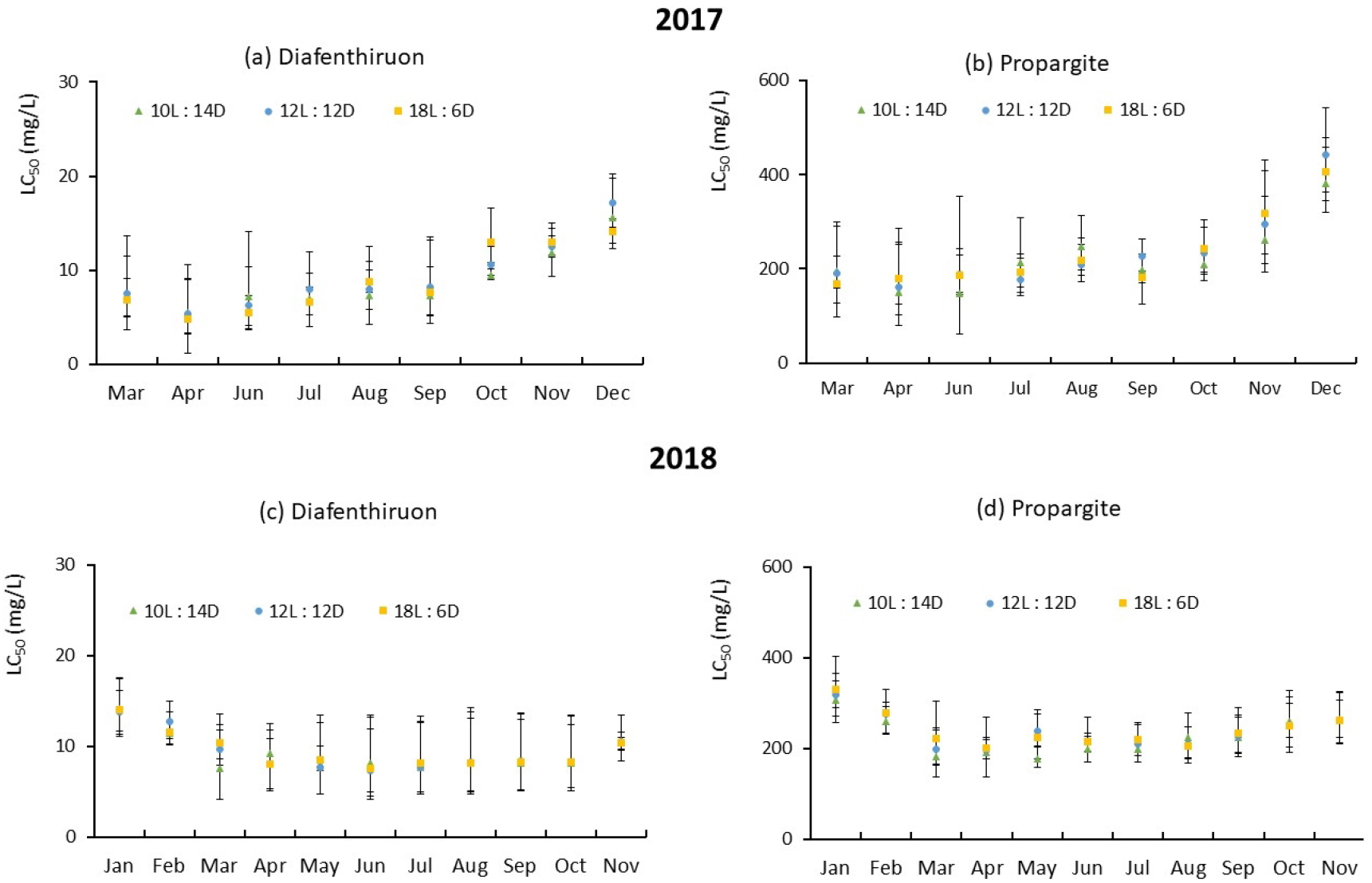
Toxicities of diafenthiruon and propargite on *T. urticae* Koch were affected by annual rhythm but not photoperiodism. The toxicities of acaricides was measured for *T. urticae* reared under three different photoperiod regimes, including 10L:14D, 12L:12D, and 18L:6D, every month from March 2017 to November 2018. The 50% lethal concentration (LC_50_ mg/L) with 95% confidence interval was used to represent the toxicity of the acaricide measured each month. The probit analyses (PROC PROBIT, SAS v9.4, SAS Institute Inc., Cary, NC, USA) was applied to understand the effects of photoperiod and time (figure a-d). The results suggested a consistent changing trend for the toxicities of both acaricides in all photoperiod regimes that the toxicity decreased in the spring and summer (from March to September) and increased in the winter (from October to Feb in the next year).

### Correlation between body size and acaricide toxicity

The body size of *T. urticae* females was measured every month from January to November 2018 to investigate potential correlation with acaricide toxicities. Our results suggested that the body size of *T. urticae* was affected by the time (F_(10, 1616)_ = 67.6, P < 0.0001), photoperiod (F_(2, 1616)_ = 11.3, P < 0.0001), and the interaction term (F_(2, 1616)_ = 8.9, P < 0.0001) (**Fig 2**). The post hoc analyses using Tukey’s HSD method suggested that *T. urticae* reared on 10L:14D has the largest body size, but the body size of the mites reared on 12L:12D and 18L:6D were not significantly different (**Table S1**). The body size of *T. urticae* reared on three photoperiod regimes varied along the time, but there was not a pattern being observed (**Fig 2**, **Table S1**). For example, the body size of *T. urticae* on the photoperiod 10L:14D had the largest body size on March, May, June, and July 2018, but lowest on September and October 2018. The size index for the treatment 18L:6D was the lowest on March, May, and July 2018. Pearson’s correlation analyses showed a weak negative correlation between the body size and the toxicities of both acaricides (**Table 1**), suggesting the smaller body size may be more sensitive to acaricides.

**Fig 2.**
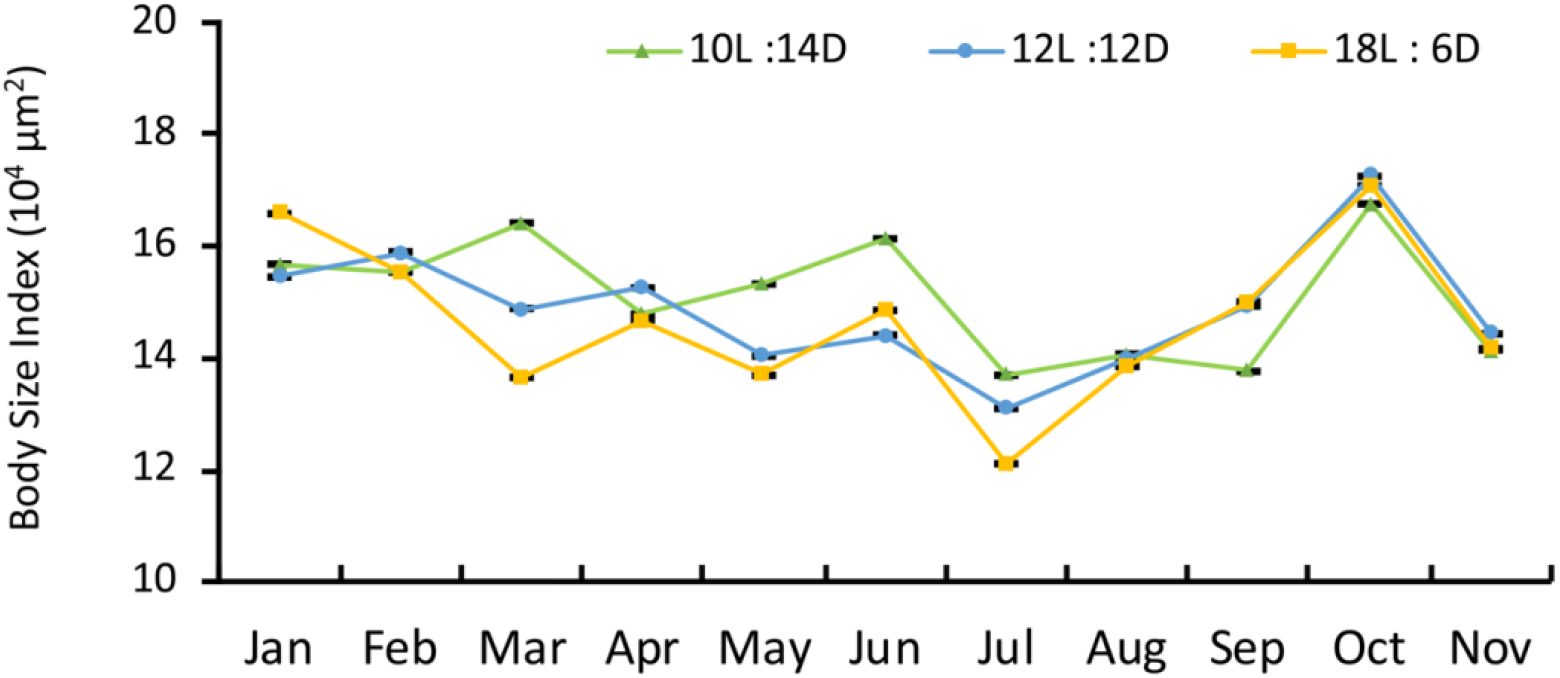
The body size of *T. urticae* is affected by both the seasonality and photoperiodism. The body size was measured every month from January to November 2018 for female *T. urticae* reared under three different photoperiod regimes, including 10L:14D, 12L:12D, and 18L:6D. A linear mixed model was applied to determine the effects of seasonality and photoperiodism. The results suggested that the body size was affected by the seasonality (F_(10, 1616)_ = 67.6, P < 0.0001), photoperiod (F_(2, 1616)_ = 11.3, P < 0.0001), and their interaction (F_(2, 1616)_ = 8.9, P < 0.0001). Tukey’s HSD method was used for post hoc comparison between different months for each of photoperiod regime and acaricide and the results are listed in Table S1. Generally, the body size reared in 10L:14D had larger body size than both in 12L:12D and 18L:6D, while the body size fluctuated over the season. The dot and error bars represent the mean and standard error for each photoperiod regime and month, respectively.

**Table 1.**
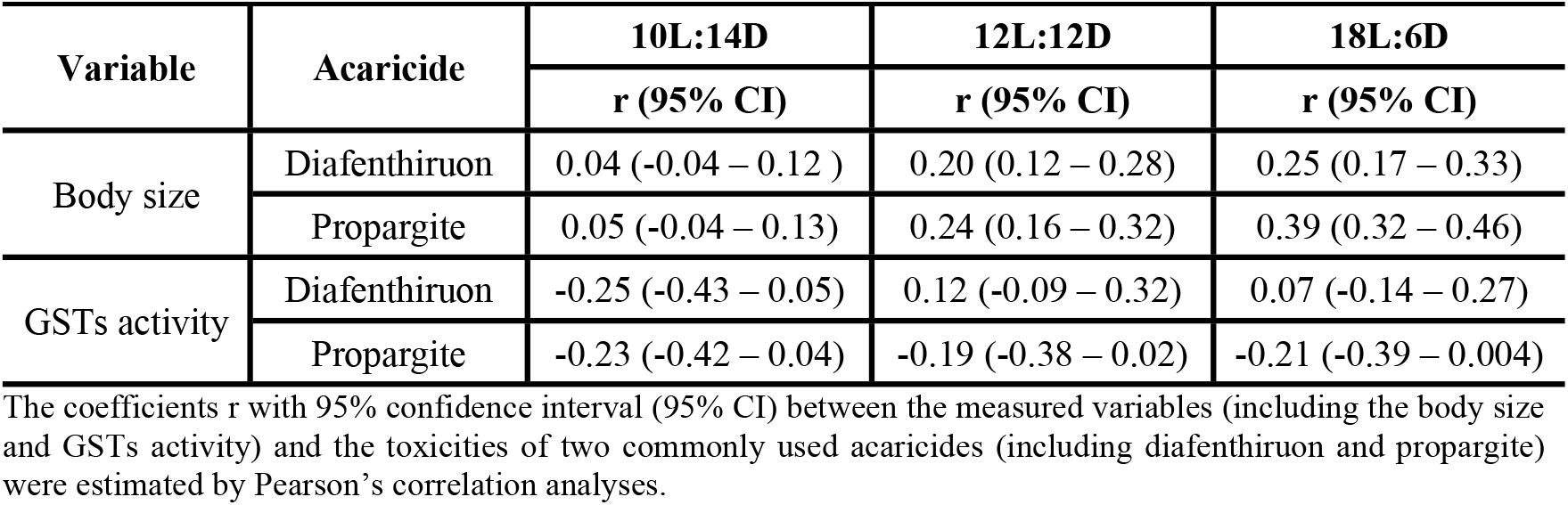
Pearson’s correlation analyses showed weak correlation between the body size and acaricide toxicities, but no correlation between the GST activities and acaricide toxicities.

### Seasonal variation of GST activities in T. urticae has no correlation with acaricide toxicity

The total GSTs activity of *T. urticae* was measured every month from January to November 2018. A linear mixed model was applied to determine the effects of time, photoperiod, and their interaction on the GST activities. The results suggested that the total GSTs activity of *T. urticae* was affected by the time (F_(10, 246)_ = 171.9, P < 0.0001), photoperiod (F_(2, 246)_ = 9.9, P < 0.0001), and the interaction term (F_(2, 246)_ = 8.6, P < 0.0001) (**Fig 3**). The post hoc analyses suggested that *T. urticae* reared on 18L:6D has the highest GSTs activity. However, the GST activities of the mites reared on 12L:12D and 14L:10D were not significantly different (**Table S2**). Along the time, there was a peak of the GSTs activity on March, 2018 for *T. urticae* reared on 18L:6D and 12L:12D, and for *T. urticae* reared on 14L:10D, the GSTs activity reached the maximum on both March and June, 2018 (**Fig 3**, **Table S2**). Pearson’s correlation analyses showed all coefficients have 95% confidence limits that overlapped with 0 (**Table 1**), suggesting no correlation between the GSTs activities and the toxicities of both acaricides.

**Fig 3.**
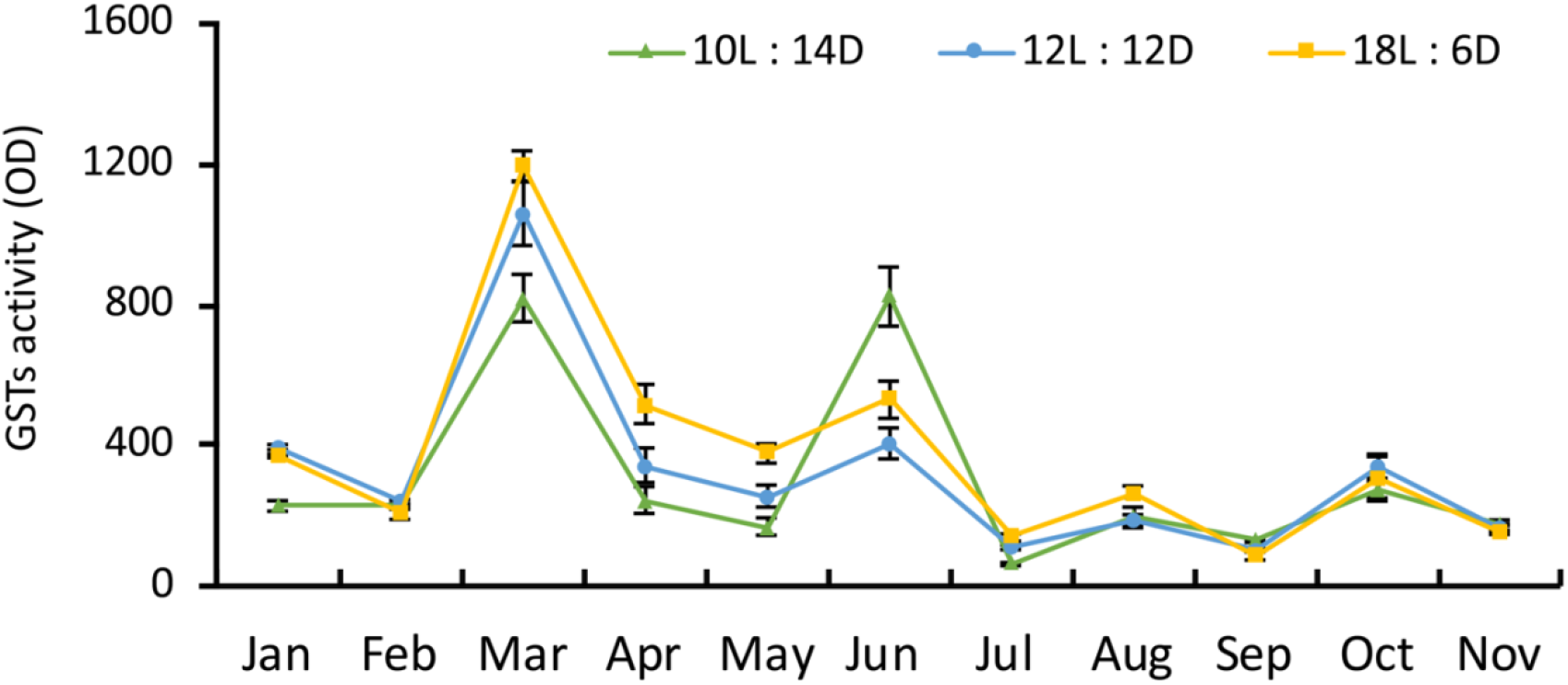
The total GSTs activity of *T. urticae* is affected by both the seasonality and photoperiodism. The total GSTs activity was measured every month from January to November 2018 for *T. urticae* reared under three different photoperiod regimes, including 10L:14D, 12L:12D, and 18L:6D. A linear mixed model was applied to understand the effects of the seasonality and photoperiodism, and results suggested that the GSTs activity was affected by the time (*F*_(10, 246)_ = 171.9, *P* < 0.0001), photoperiod (*F*_(2, 246)_ = 9.9, *P* < 0.0001), and the interaction term (*F*_(2, 246)_ = 8.6, *P* < 0.0001). Tukey’s HSD method was used for *post hoc* comparison between different months for each of photoperiod regime and acaricide and the results are listed in Table S2. The dot and error bars represent the mean and standard deviation for each photoperiod regime and month, respectively.

## Discussion

In this study, we investigated the effects of seasonality and photoperiodism on the toxicities of two acaricides to control the two-spotted spider mite, *T. urticae.* Our results showed that acaricide toxicities were not significantly different among the three photoperiods (10L:14D, 12L:12D and 18L:6D). However, toxicities changed consistently across the seasons even though other abiotic factors including photoperiod, temperature, and relative humidity were controlled. Toxicities increased in spring and summer (from March to September) and decreased in the winter (from October to February in the next year) in all photoperiod regimes. We measured the body size and total GSTs activity every month from January to November in 2018 to determine their correlation to the toxicities. Even though the body size and total GSTs activity were found to be affected by both photoperiodism and seasonality, there was only a weak negative correlation between the body size and the toxicities, and no correlation between the total GSTs activity and the toxicities.

Our results showed that the photoperiodism affects physiology of *T. urticae* including the body size and total GSTs activity, but not the pesticide toxicity. This is consistent with our previous study which only examined the acaricide toxicity at a single time point in the summer of 2016 [19]. Photoperiodism, reflecting the daily variation of day length, plays a crucial role in modulating physiological changes in different arthropod species, for example, diapause, induction of seasonal morphs, and immature development [2, 4]. Our results are consistent with this showing the photoperiodism has strong effects in body size and GSTs activities of *T. urticae*. The cascading changes in the physiology have been proposed to alter the sensitivities of arthropod pests against pesticides. For example, the altered photoperiods could induce diapause and several key physiological and genetic changes such as metabolic suppression and regulation of enzymatic activities, further influencing the capabilities of insects detoxifying pesticides [26-28]. A short day time (8L:16D) induced the up-regulated detoxification enzymes including esterase, GSTs and P450s in a lepidopteran pest, *Spodoptera litura* (Lepidoptera: Noctuidae) [29]. In this study, instead of measuring the toxicities at specific time point, we extended the understanding about the effects of photoperiodism on the acaricide toxicities on *T. urticae* to a continuous period of two consecutive years, and found the photoperiod regime and its interaction with the time/season did not have significant effects on the toxicities.

Contrary to the effects of photoperiodism, the toxicities of both acaricides on *T. urticae* changed across the seasons. Seasonality has been associated with diverse abiotic factors and have been suggested to alter the physiology and pesticide toxicity when measured in the field-collected populations [12, 13]. The seasonal variation of resistance to different kinds of pesticides in insects varies in the field. For example, a 6-year investigation to the effects of seasonal changes on the level of resistance of *Helicoverpa armigera* to pesticides indicated the level of resistance to quinalphos and endosulfan increases in the winter (late November), but to pyrethroids be lower from mid-October to early November compared with seasonal average [30]. A recent study of seasonally changed pesticide toxicity in the field populations of *Anopheles hyrcanus* and *Culex pipiens* complex mosquitoes showed the seasonal cycling pattern of pyrethroid and organophosphate toxicities partially resulted from the altered frequency of resistant mutation across the season [31]. However, in field-sampled experiments, effects from other changing abiotic factors cannot be excluded. In this study, we reared *T. urticae* in incubators with the temperature, relative humidity, and light intensity kept consistent for all three photoperiod treatments. Our results showed an intrinsic rhythm changing trend over the season, which is also not affected by the photoperiodism. While we seek to determine if body size and detoxification metabolize enzymes such as the GSTs activity can affect acaricide toxicity we only found a weak and negative correlation between the acaricide toxicity and body size. Body size is a trait involving complex genetic effects, of which the addition of both positive and negative contributions could compositely lead to the weak negative correlation.

Further understanding of circadian or seasonal variation underlying acaricide toxicity in *T. urticae* is useful to inform the management strategies such as by better timing of pesticide application, which can reduce the use of pesticides and delay the occurrence and development of pesticide resistance in the spider mite [2, 7, 30]. In this multi-year study, the toxicities of diafenthiruon and propargite to T. urticae were found modulated by seasonality, decreasing in the winter and increasing in the summer, but not affected by photoperiodism. Therefore, in the field, except for considering the effects of abiotic factors in different seasons, the acaricide timing and dosage should be adjusted to coordinate with the endogenously changing toxicities, which can be beneficial by preventing the development of acaricide resistance in the next season [32, 33]. In addition, optimal timing of pesticide application based on understanding the endogenous effects on the pesticide toxicity can also be applicable to other pest species [12, 13, 34]. These considerations in pest management strategies can improve the efficacy of chemical controls and contribute to a safer environment.

## Conclusions

In conclusion, we used a worldwide agricultural pest, *T. urticae*, as a model to investigate how photoperiodism and seasonality affect pesticide toxicity. The results showed that seasonality but not photoperiodism affects pesticide toxicity of propargite and diafenthiruon on *T. urticae*. The toxicities of both pesticides were found not significantly different among the three photoperiod regimes. However, the toxicities were found cyclically changing over the season which increased in the summer and decreased in the winter. We found photoperiodism and seasonality had complex effects on the body size and GSTs activity, but only the body size had a weak negative correlation with the pesticide toxicities. Our study is the first to show that seasonality has an endogenous effect on the pesticide toxicity, which is important for pest control practice in the field.

## Acknowledgments

The authors would like to thank Dr. Wenxin Liu and Xingxing Liu (College of Agronomy and Biotechnology, China Agriculture University, China) for helping us with data analyses. Y.L. acknowledges the funding support from the National Natural Science Foundation of China (31560520), Yunnan Science and Technology Foundation of China (2017HB059) and SAFEA's Overseas Training of China (P182039008).

## Supporting information

**S1 Fig. Toxicities of diafenthiruon and propargite on *Tetranychus urticae* Koch with each photoperiod regime.**

**S1 Table. Post hoc analyses of *T. urticae* body size among photoperiod regimes, months, and the interactions of photoperiod regime and month, respectively**

**S2 Table. Post hoc analyses of *T. urticae* GSTs activity among photoperiod regimes, months, and the interactions of photoperiod regime and month, respectively.**

## Author Contributions

**Conceptualization:** Yanjie Luo and Henry Chung.

**Data curation:** Zhenguo Yang and Zinan Wang.

**Formal analysis:** Zhenguo Yang and Zinan Wang.

**Funding acquisition and project administration:** Yanjie Luo.

**Investigation and laboratory work:** Zhenguo Yang, Jing Ni, Aisi Da and Daoyan Xie.

**Writing—original draft:** Yanjie Luo and and Zinan Wang.

**Writing—review and editing:** Henry Chung.

**All authors read, corrected, and approved the manuscript.**

